# Calorie Restriction-Induced Daily Hibernation in Mice Drives Cyclic DNA Damage and Repair

**DOI:** 10.64898/2026.06.02.729536

**Authors:** Fia Fia Lie, Maurits Roorda, Maaike Goris, Femke Hoogstra-Berends, Roelof A. Hut, Marco Demaria, Robert H. Henning

**Affiliations:** Department of Clinical Pharmacy and Pharmacology, University of Groningen, University Medical Center Groningen, Netherlands; Department of Pharmacology, Faculty of Medicine, Universitas Tarumanagara, Jakarta, Indonesia; Chronobiology unit, Groningen Institute of Evolutionary Life Sciences, University of Groningen, Netherlands; European Research Institute for the Biology of Ageing (ERIBA), University of Groningen, University Medical Center Groningen, Netherlands

**Keywords:** Calorie restriction, hibernation, DNA repair, metabolic challenge, genomic stability

## Abstract

Hibernation consists of bouts of torpor, characterized by profound decreases in metabolism and body temperature (T_b_), alternated with periods of euthermia called interbout arousals, during which normal metabolism and T_b_ resume. Seasonal hibernators accumulate DNA strand breaks during torpor, which are repaired during arousal. Here, we assess dynamics of DNA damage and repair during serial daily torpor in mice induced by 30% calorie restriction (CR) and investigate the effects of metabolic challenge on DNA repair. Serial daily torpor induced by CR in C57/BL6J mice of both sexes housed at 20°C lasts 6–12 hours. Like seasonal hibernators, DNA damage increases in CR-induced torpor and is repaired in the subsequent euthermic period, as evidenced by comet assay and ɣH2AX accumulation. To metabolically challenge animals, ambient temperature (T_a_) was lowered to 4°C, since torpid mice defend a T_b_ of around 20°C or higher. Despite inducing a significant metabolic challenge, housing of torpid mice at 4°C does not increase DNA damage compared to 20°C housing. However, reducing T_a_ to 4°C during euthermia inhibits DNA repair. Interestingly, p21 levels increase in mice exposed to 4°C, indicating cell-cycle inhibition during exposure to 4°C. Thus, 30% CR induces daily cycles of torpor-induced DNA damage and euthermia-associated DNA repair in mice, and exposure to a T_a_ of 4°C during arousal inhibits DNA repair mounting a cell cycle inhibition response. Thus, the torpor-arousal cycle may be a contributing factor to the lifespan extension benefits of CR in mice, promoting genomic integrity and thereby cellular and tissue health.

## Introduction

Hibernation is an adaptive mechanism for coping with unfavorable environmental conditions. Throughout hibernation, seasonal deep hibernators alternate multi-day torpor bouts with shorter periods of euthermic arousal. Torpor bouts are marked by significant decreases in metabolism, body temperature (T_b_), heart rate, respiratory rate, and other physiological parameters, whereas in arousals metabolism and physiology return to normal (1). Torpor bouts typically last from several days to weeks while arousal mostly lasts less than a day. Other animals, particularly smaller mammals, may enter torpor on a daily basis, known as daily torpor (2). It is assumed that similar molecular mechanisms underlie multi-day and daily torpor, as multiple species employ both (3, 4).

While the torpor phase offers substantial energy-saving benefits, it is also marked by the molecular signatures of cellular damage, such as tau hyperphosphorylation in neurons (5–7), extracellular matrix deposition in lung (8) and activation of autophagy in heart (9). Remarkably, this torpor associated damage is reversible and fully resolved during the subsequent arousal, indicating rapid and substantial repair. Recently, we documented the accumulation of DNA strand breaks during torpor in the seasonal hibernator, garden dormouse (*Eliomys quercinus*) (10). Consistent with this observation, pathways related to DNA damage, cell cycle arrest, and cell death are prominently regulated in the liver transcriptome of hibernating golden hamsters (*Mesocricetus auratus*) (11). To date, it is unknown whether torpor-induced DNA damage also occurs in daily hibernation in mice which is characterized by much shorter torpor phases at a higher T_b_ of ∼20°C.

Additionally, the origin of torpor-induced DNA damage remains unclear. Two main hypotheses were proposed. The first comprises an increase in the oxygen radical load because of the suppression of metabolism and lowering of T_b_ during torpor, as documented in cooled cells (12). Alternatively, inhibition of the repair of stochastic DNA damage may underlie strand break accumulation during torpor. To study these hypotheses, we used C57/BL6J mice (*Mus musculus*) implanted with a temperature logger, in which serial daily torpor was induced by 30% calorie restriction (CR) (13, 14) and quantified T_b_, metabolism, and DNA damage. In addition, given that torpid mice defend a T_b_ of around 20°C, we metabolically challenged mice during specific phases by housing them at low ambient temperature (T_a_ 4°C) during torpor, arousal, or both. Our results document the accumulation of DNA damage in daily torpor in calorie restricted mice, and this DNA damage is repaired during the subsequent arousal. We find that decreasing the ambient temperature during arousal, but not during torpor, exacerbates DNA damage. Thus, whereas a metabolic challenge during torpor does not increase DNA damage, we postulate it halts DNA repair during euthermia.

## Results

### Calorie restriction induces daily torpor, and lowering T_a_ decreases T_b_ and increases metabolic rate

Mice housed at 20°C were either fed ad libitum (adlib) or subjected to 30% CR by feeding at zeitgeber (ZT) 8 to induce daily torpor. A metabolic challenge was inflicted upon designated groups by lowering the T_a_ to 4°C during their last torpor bout and/or during their subsequent final euthermic period (henceforth indicated as ‘arousal’). Mice were euthanized either during a torpor bout (T, ZT 23) or during late arousal (A, ZT 17). Consequently, mice were divided into eight experimental groups: Adlib20, Adlib4, and six CR groups: T20, T4, T20A20, T20A4, T4A20, and T4A4 (Fig. 1A). Torpor was defined as a decrease in T_b_ ≥5°C (2), and periods with T_b_ ≤31°C were classified as torpor and T_b_ >31°C as arousal.

**Figure 1.**
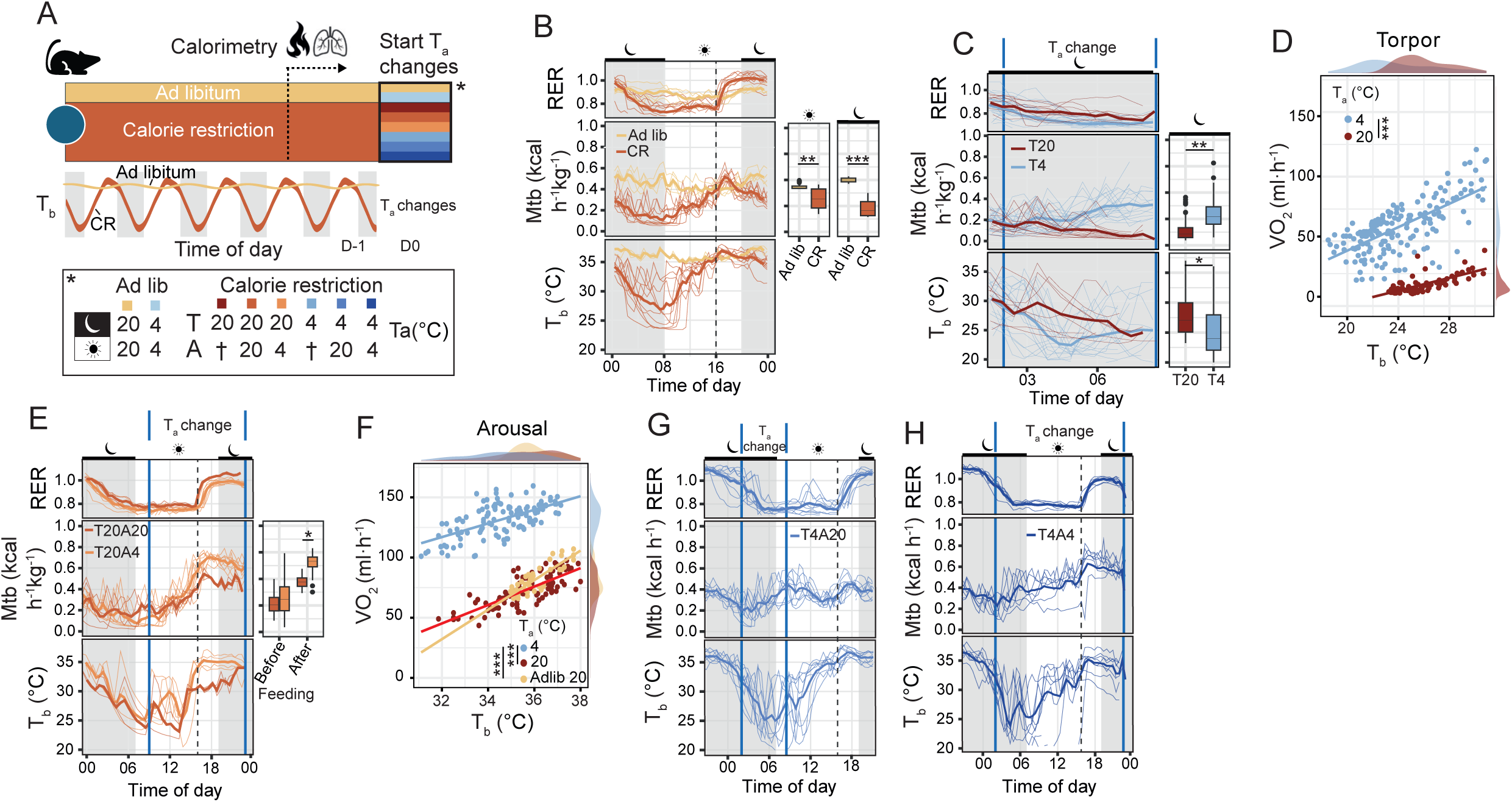
Calorie restriction induces daily hibernation, and lowering T_a_ decreases T_b_ and increases metabolic rate. **(A)** Experimental timeline: Mice were implanted with temperature loggers and allowed 1 week recovery. Following 1 week recovery, mice were randomized to eight groups and were provided food ad libitum (2 groups) or gradually subjected to 30% calorie restriction over two weeks (6 groups). The six groups on 30% calorie restriction, displayed consistent serial daily hibernation, defined by daily torpor and arousal. To inflict a metabolic challenge, T_a_ was changed on the last day of the experiment (D0) followed by sacrifice. Two ad libitum fed groups were housed at constant T_a_ of 20°C (adlib20, n = 4) or subjected to T_a_ of 4°C during the time period corresponding to torpor (20:00 to 08:00, adlib4, n = 4). Two groups were housed at constant T_a_ of 20°C and sacrificed after torpor (T20, n = 12) or arousal (T20A20, n = 4). Two groups were housed at constant T_a_ of 4°C and sacrificed after torpor (T4, n = 24) or after arousal (T4A4, n = 9). Two groups were housed alternating T_a_ between 20°C and 4°C during torpor and arousal and sacrificed after arousal (T4A20, n = 9, and T20A4, n = 8). T=torpor; A=arousal (arousal); 20=20°C; 4=4°C. **(B)** Respiratory exchange ratio (RER), metabolic rate and T_b_ of ad libitum and CR animals exhibiting daily cycles of torpor and arousal. Light-dark cycles are indicated as shaded areas, feeding times (16:00) are indicated with dashed lines. Data are represented as median and interquartile ranges. **(C)** RER, metabolic rate and T_b_ of torpid animals exposed to T_a_ of 20°C and 4°C between 02:00 and 08:00. n_T20_ = 8, n_T4_ =24 **(D)** T_b_ versus VO2 consumption of torpid animals exposed to T_a_ of 20°C and 4°C. n_T20_ = 8, n_T4_ =24. Data are represented as individual measurements of animals with a linear fit. **(E)** RER, metabolic rate and T_b_ of arousing animals exposed to T_a_ of 20°C and 4°C between 08:00 and 20:00. n_T20A20_ = 4, n_T20A4_ = 8. **(F)** T_b_ versus VO2 consumption of arousing animals exposed to T_a_ of 20°C and 4°C, and ad libitum animals exposed to T_a_ of 20°C. n_adlib20_ = 4, n_T20A20_ = 4, n_T20A4_ = 8. Data are individual measurements of animals with a linear fit. **(G)** RER, metabolic rate and T_b_ of torpid animals exposed to 4°C between 02:00 and 08:00, followed by arousal at T_a_ of 20°C n_T4A20_ = 9. **(H)** RER, metabolic rate and T_b_ of animals exposed to T_a_ of 4°C during both torpor and arousal. n_T4A4_ = 9. *P<0.05, **P≤0.005, ****P≤0.0001.

*Ad libitum* fed animals showed the expected fluctuations in daily T_b_ and metabolism according to the light:dark cycle (Fig. 1B). CR at 30% robustly induced bouts of daily torpor (Fig. 1B, Table 1, Supp. Fig1A-D). T_b_ during torpor bouts reached minima of 22°C, with gradual restoration of T_b_ to euthermia upon arousal (Fig. 1B), despite significant variance between animals in torpor depth and patterns. Upon feeding, CR mice showed a respiratory exchange ratio (RER) of 1.0, indicative of carbohydrate combustion, followed by a gradual decline towards fatty acid oxidation (RER ≈ 0.75) during the second half of the dark/torpid phase until feeding on the subsequent day (Fig. 1B). Total metabolism was markedly reduced in CR compared to ad libitum feeding across the light:dark cycle (Fig. 1B). Further, upon the initiation of a torpor bout, metabolic rate of CR mice decreased faster than T_b_, consistent with active hypometabolism similar to observations in seasonal hibernators (Supp. Fig. 1E). Taken together, CR mice underwent robust serial daily torpor characterized by lipid combustion and a drastic decrease in metabolism and T_b_.

**Table 1.** Metabolism and body temperature (T_b_) measurements by indirect calorimetry.

| Group | Metabolism torpor (kcal/h.kg) | Metabolism arousal (kcal/h.kg) | $T_b$ torpor ($^{\circ}\text{C}$) | $T_b$ arousal ($^{\circ}\text{C}$) |
| --- | --- | --- | --- | --- |
| Adlib20 | $0.47 \pm 0.05$ (n=4) | $0.47 \pm 0.03$ (n=4) | $36.8 \pm 0.8$ (n=2) | $36.8 \pm 0.8$ (n=2) |
| Adlib4 | $0.67 \pm 0.08$ (n=4) | n.d. | $35.2 \pm 0.8$ (n=2) | n.d. |
| T20 | $0.11 \pm 0.02$ (n=12) | $0.29 \pm 0.03$ (n=6) | $27.1 \pm 1.5$ (n=8) | $33.6 \pm 0.6$ (n=6) |
| T4 | $0.24 \pm 0.03$ (n=19) | $0.31 \pm 0.08$ (n=19) | $25 \pm 1.1$ (n=19) | $32.9 \pm 1.1$ (n=19) |
| T20A20 | $0.11 \pm 0.01$ (n=4) | $0.35 \pm 0.07$ (n=4) | $26.8 \pm 0.2$ (n=2) | $32.7 \pm 0.1$ (n=2) |
| T20A4 | $0.12 \pm 0.04$ (n=4) | $0.52 \pm 0.04$ (n=4) | $26.7 \pm 1.0$ (n=4) | $34.2 \pm 0.7$ (n=4) |
| T4A20 | $0.25 \pm 0.02$ (n=5) | $0.34 \pm 0.04$ (n=5) | $26.4 \pm 1.5$ (n=5) | $34.8 \pm 0.7$ (n=5) |
| T4A4 | $0.32 \pm 0.00$ (n=3) | $0.55 \pm 0.01$ (n=3) | $26.3 \pm 0.4$ (n=3) | $33.7 \pm 1.3$ (n=3) |

Data are presented as mean ± SD; n= number of animals included in the analysis. N.d. = not determined as these groups were only cooled during the time corresponding to torpor.

To inflict a metabolic challenge, T_a_ was lowered to 4°C, which increased metabolism and decreased body temperature in ad libitum fed animals (Supp Fig. 1C, F). In CR groups, T_a_ was lowered to 4°C either during torpor (T4), arousal (T20A4), or throughout both phases (T4A4). Lowering T_a_ during torpor (T4) significantly increased metabolism compared to normal torpor (T20), while reducing T_b_ (Fig. 1C). T4 mice exhibited higher oxygen consumption (VO2) compared to T20, even at the same body temperature (Fig 1D). Lowering T_a_ during arousal (T20A4) significantly increased metabolism compared to normal arousal (T20A20), especially after feeding (Fig. 1E). Similarly, T20A4 mice exhibited higher VO2 compared to T20A20 at the same T_b_ (Fig. 1F).

In addition to affecting metabolism and T_b_, low T_a_ also disturbed torpor patterns. First, lowering of T_a_ during torpor instigated inconsistent torpor in some animals, reflected by multiple brief arousals (Fig. 1G), likely reflecting the need to regularly increase metabolism to defend T_b_. Furthermore, lowering of T_a_ during arousal (T20A4, T4A4) prolonged torpor duration compared to the previous day (Supp. Fig. 1G), as demonstrated by continued metabolic depression and lower T_b_ until feeding at 4PM (Fig. 1H).

Overall, CR mice underwent serial daily torpor, with a significant reduction in metabolism and T_b_. Lowering T_a_ to 4°C during torpor and arousal resulted in a higher metabolic rate, lower T_b_ during torpor, and extension of torpor duration.

### Calorie restriction induces daily cycles of DNA damage during torpor and repair during arousal, repair is inhibited by metabolic challenge

To explore DNA damage and repair, we quantified total DNA strand breaks (without distinction between single and double strand breaks) and double-strand DNA breaks (DSBs) using alkaline and neutral comet assays on freshly isolated splenocytes. Ad libitum fed mice showed low amounts of basal DSBs and SSBs, which were unaffected by housing them at a T_a_ of 4°C (Fig. 2A-B, Supp. Fig. 2A). In CR mice, DSBs accumulated during torpor, which were resolved in the subsequent arousal indicating DSB repair (Fig. 2A-B). Similarly, in the alkaline comet assay that quantifies both SSBs and DSBs, we observed a small but significant increase in DNA damage at the end of torpor, followed by repair during arousal (Supp. Fig. 2A). Variance in DNA damage between animals in the same group was considerable (Supp. Fig. 2B), which probably relates to the substantial variance in T_b_, metabolism and torpor bout duration of the CR model (Fig. 1B, Supp. Fig. 1G). Investigation of the relationship between mean DNA damage per animal (T20 and T4) and age, metabolic rate during torpor and mean T_b_ during torpor did not reveal any significant interactions, thus leaving the variance observed between animals unexplained (Supp. Fig. 1C).

**Figure 2.**
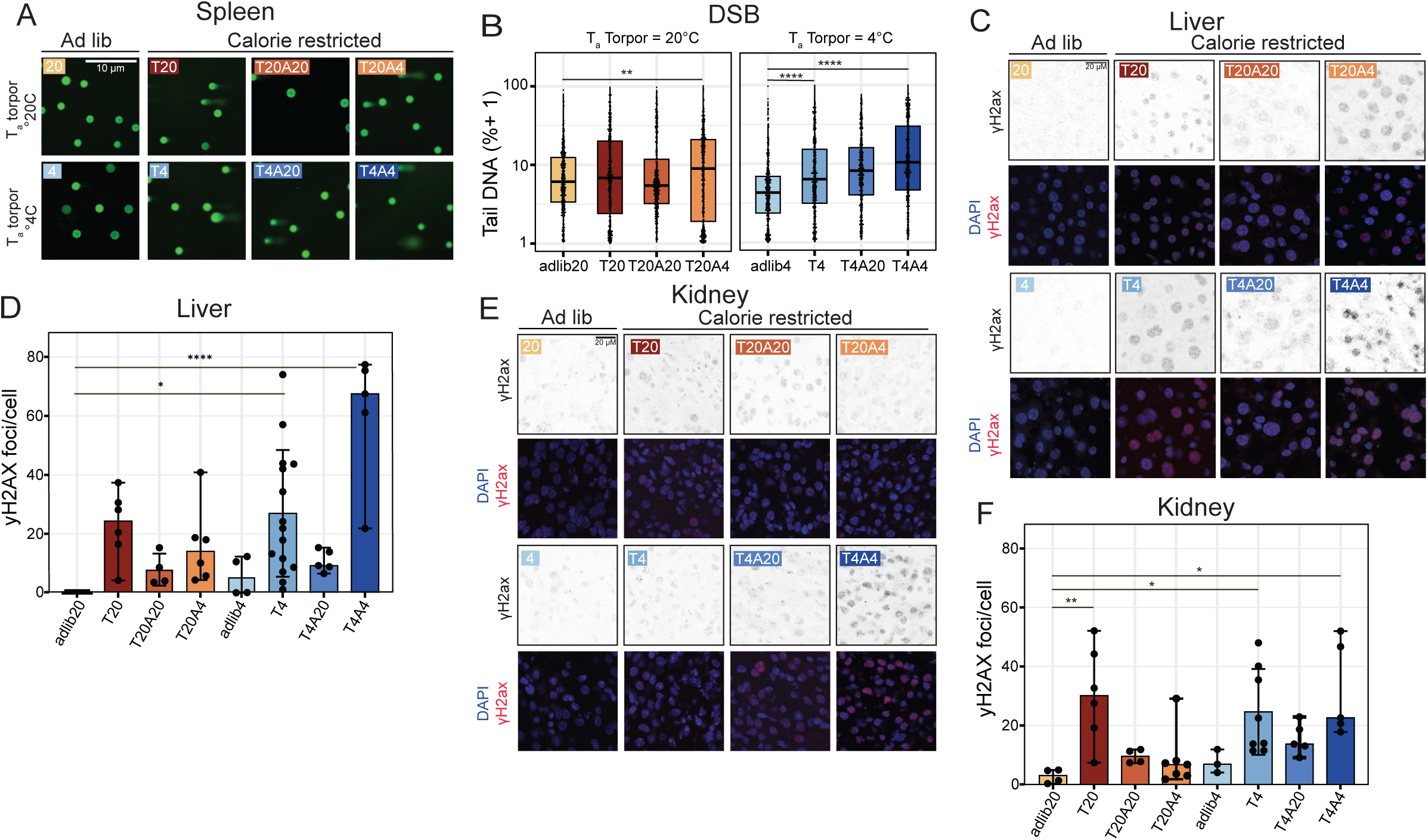
Calorie restriction induces daily cycles of DNA damage during torpor and repair during arousal, and repair is inhibited by metabolic challenge. **(A)** Representative pictures from alkaline comet assays performed on freshly isolated splenocytes from animals in all groups indicated. Scale bar: 10µM. **(B)** Quantification of neutral comets (DSB only) of ad libitum fed and CR animals housed at different T_a_. Data are represented as median and interquartile ranges. **(C)** Representative images of ɣH2AX immunostaining in liver of ad libitum fed and CR animals housed at different T_a_. Scale bar: 20µM. **(D)** Quantification of liver ɣH2AX foci from (C) presented as mean of foci/cell per animal. Data are represented as mean ± SD **(E)**Representative images of ɣH2AX immunostaining in kidney of ad libitum fed and CR animals housed at different T_a_. Scale bar: 20µM. **(F)** Quantification of liver ɣH2AX foci from (E) presented as mean of foci/cell per animal. Data are represented as mean ± SD. *P<0.05, **P≤0.005, ****P≤0.0001.

Next, we explored whether a metabolic challenge during torpor or arousal affected DNA strand breaks. During torpor at low T_a_ (4°C), total DSBs accumulated similarly in T4 compared to T20 animals (Fig. 2A-B, Supp. Fig 2A), indicating that a higher metabolic rate during torpor does not induce excess DNA damage. We then examined the effects of metabolic challenges during arousal. Mice exposed to low T_a_ solely during arousal (T20A4) retain residual DSBs compared with Adlib20 (Fig. 2A-B, Supp. Fig 2A). Importantly, increased DNA strand breaks also persisted in the arousal of mice facing a metabolic challenge during both torpor and arousal (T4A4), which showed significantly higher total strand breaks and DSB (Fig. 2A-B, Supp. Fig 2A). To further substantiate that torpor induced DNA strand breaks in CR mice, we measured levels of phosphorylated histone 2AX (ɣH2AX) in liver and kidney, two major metabolic organs. In line with our observations in spleen, we observed an increase in ɣH2AX foci during torpor, which resolve upon arousal both in liver (Fig 2C-D) and kidney (Fig. 2E-F). In both liver and kidney, lowering T_a_ during torpor to 4°C did not produce further ɣH2AX foci, but lowering T_a_ during arousal did (Fig. 2C-F).

Overall, these observations demonstrate that CR induces daily cycles of DNA damage accumulation in torpor and repair during arousal in multiple tissues. Further, DNA repair is inhibited by metabolically challenging the animals by lowering T_a_ during arousal, possibly even exacerbating DNA damage.

### Arousal at low T_a_ triggers sustained DNA damage signaling and cell cycle repression by p21

Next, we explored the consequences of the torpor-induced DNA damage and repair in the liver and kidney. We quantified protein levels and mRNA expression of p21, a potent inhibitor of cyclin-dependent kinases, which is activated by multiple stressors including DNA double-stand breaks, and which inhibits cell cycle progression but permits DNA repair (15, 16).

In *ad libitum* fed animals, lowering T_a_ to 4°C increased liver p21 protein expression (Adlib20 vs. Adlib4, Fig. 3A-B). In 30% CR, p21 protein levels decreased during torpor and restored during arousal, with no effect of lowering T_a_ during arousal (Adlib20 vs. T20, T20 vs. T20A20 and T20A4, Fig. 3A-B). Lowering T_a_ to 4°C during torpor increased p21 protein levels, which are fully rescued by arousing at T_a_ of 20°C. In contrast, during arousal at T_a_ of 4°C p21 protein levels remain high (T20 vs. T4, T4 vs. T4A20 and T4A4, Fig 3B), indicating that cell cycle inhibition by p21 in the liver is relieved during torpor, but reinstituted at a T_a_ of 4°C during torpor (T4) or torpor and arousal (T4A4). Since p21 is a core target of p53 upon sensing DNA damage, we investigated its mRNA expression using RNAscope and qPCR as a proxy for p53 activity. The mRNA expression of p21 in liver significantly increased in torpid mice maintained at 4°C during torpor (adlib20 vs. T20A4 and T4, Fig. 3C-E). This increase persisted in mice exposed to cold conditions during torpor and arousal (T20 vs. T4A4, Fig. 3C-E). In contrast, returning mice to 20°C during arousal significantly decreased p21 mRNA expression (T4 vs. T4A20 and T4A20 vs. T4A4, Fig. 3C-E). Expression of p21 mRNA in kidney tissue as measured by RNAscope showed similar patterns to those observed in liver, but with lower p21 expression in general. p21 mRNA increased expression during torpor at a T_a_ of 4°C, which persisted in mice exposed to T_a_ of 4°C during torpor and arousal (Fig. 3F-G). Overall, these observations demonstrate that CR induces daily cycles of reversible modulation of cell-cycle inhibitory signaling. Further, infliction of a metabolic challenge with consequent increase in DNA damage increased p21 protein levels and expression during cold torpor, which remained high during cold arousal. This indicates lowering T_a_ during torpor induces cellular stress and likely activates p21 through DNA damage signaling, leading to cell cycle inhibition signaling upon metabolic challenge.

**Figure 3.**
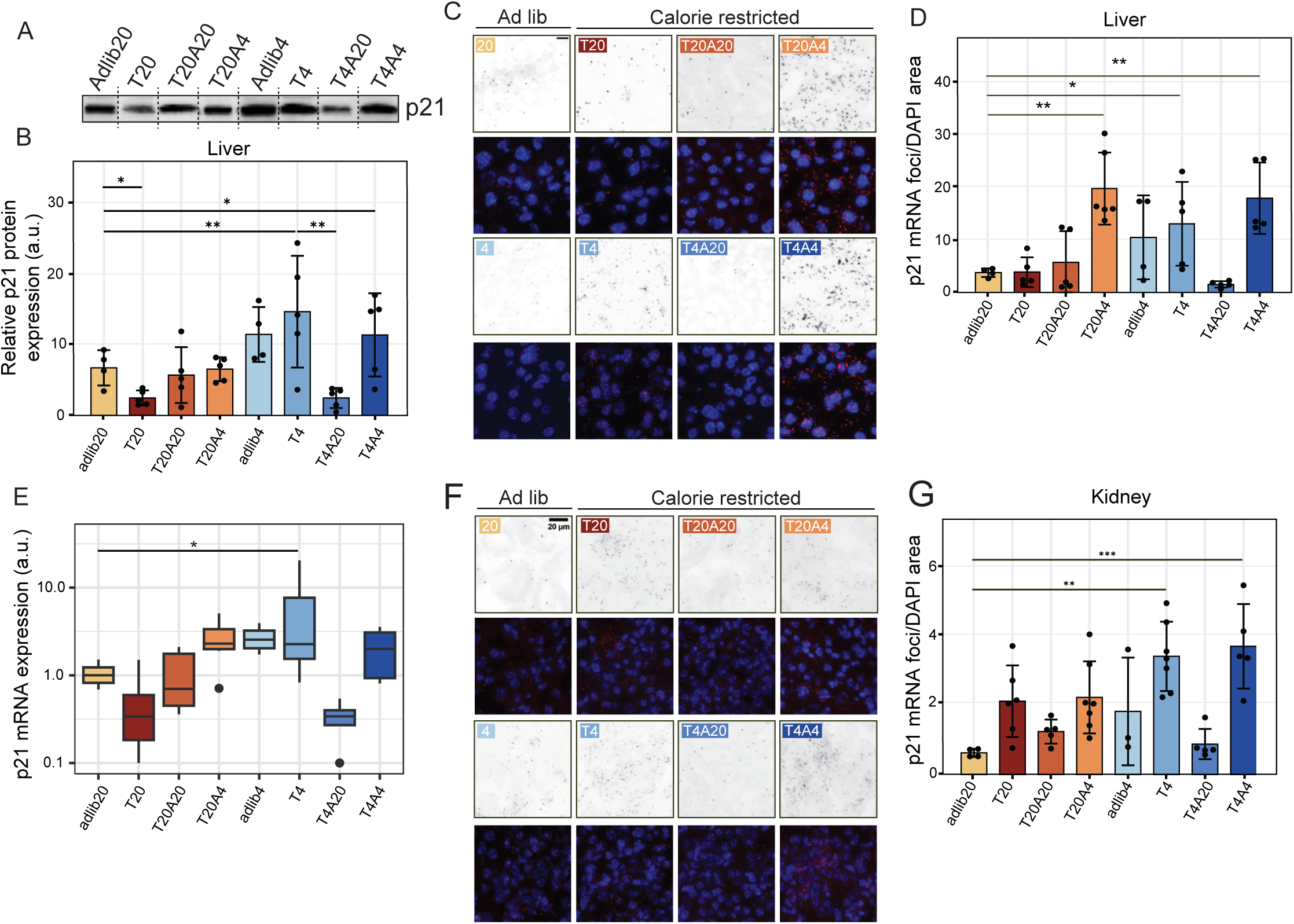
Arousal at low T_a_ triggers sustained DNA damage signalling and cell cycle repression by p21. **(A)** Representative western blot images of p21 protein in liver of ad libitum fed and CR animals housed at different T_a_. **(B)** Quantification of p21 protein levels shown in (A). Data are represented as mean ± SD **(C)** Representative images of p21 mRNA foci in liver. Scale bar: 20µM. **(D)** Quantification of liver p21 mRNA foci from (C) presented as mean of foci/DAPI area per animal. Data are represented as mean ± SD **(E)** Quantification of liver p21 mRNA expression using qPCR of ad libitum fed and CR animals housed at different T_a_. Data are represented as mean ± SD **(F)** Representative images of p21 mRNA foci in kidney. Scale bar: 20µM. **(G)** Quantification of kidney p21 mRNA foci from (F) presented as mean of foci/DAPI area per animal. Data are represented as mean ± SD. *P<0.05, **P≤0.005, ****P≤0.0001.

### Calorie restriction induces a specific antioxidant and metabolic gene expression program, putting animals in cold arousal at risk of ROS formation

Our observation that metabolism drops more sharply than body temperature while entering torpor led us to hypothesize that torpor, arousal and changing T_a_ will influence expression of genes shaping the response to changes in metabolism. Therefore, we measured mRNA expression of nuclear factor erythroid 2-related factor (Nrf2), hypoxia inducible factor 1 alpha (Hif1a) and manganese superoxide dismutase (mnSOD), all important players in antioxidant responses associated with changes in energy source and metabolic rate. Nrf2 transcription decreased in all CR groups, but notably also in ad libitum fed animals at T_a_ of 4°C (Fig. 4A). Arousal at T_a_ of 20°C increased Nrf2 transcription, but arousal at T_a_ of 4°C did not (A20 groups vs. A4 groups, Fig. 4A). HIF-1a transcription decreased in animals that spent torpor and arousal at 4°C (adlib4 vs T4A4, Fig. 4B). mnSOD expression transcription was decreased in mice exposed to 4°C throughout torpor and arousal (adlib4 vs. T4A4, Fig. 4C). Thus, mRNA expression of Nrf2 significantly decreased in CR groups, and HIF-1a and MnSOD were markedly lower in mice exposed to 4°C during torpor and arousal (T4A4). Further, the relation between Nrf2 transcription and metabolic rate seems inverted in metabolically challenged mice: placement at a T_a_ of 4°C increased metabolic rate yet decreased Nrf2 transcription. Together, this could indicate that CR drives a specific antioxidant gene expression program that is diametrically opposed to expression of the same genes under ad libitum fed conditions.

**Figure 4.**
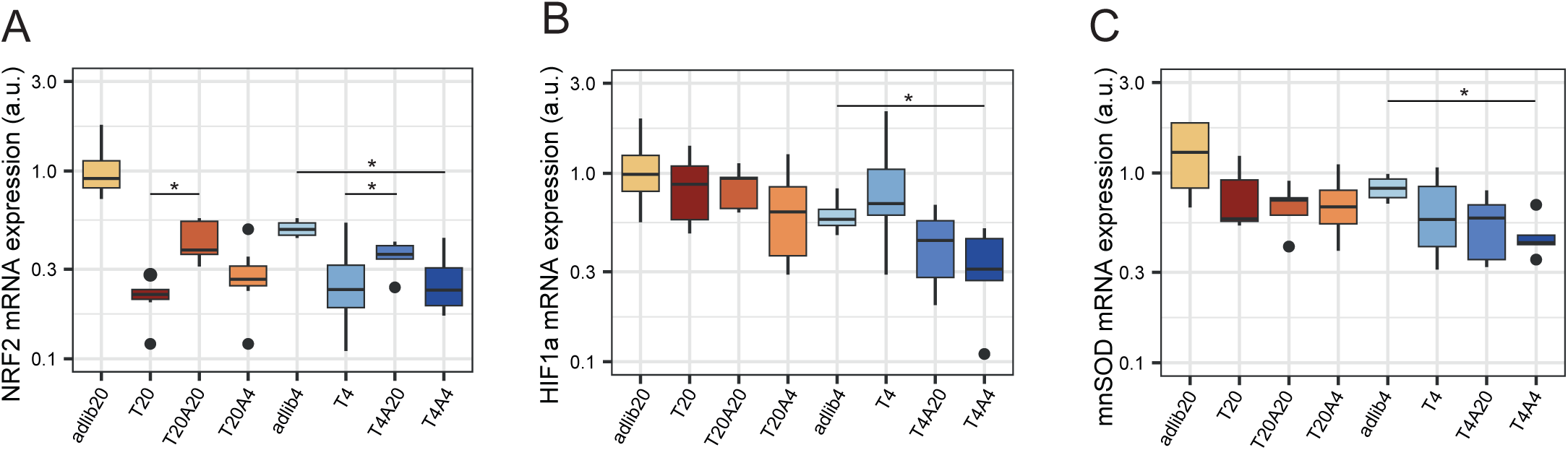
Calorie restriction induces a specific antioxidant and metabolic gene expression program, putting animals in cold arousal at risk of ROS formation. **(A)** Quantification of liver NRF2 mRNA expression using qPCR of ad libitum fed and CR animals housed at different T_a_. Data are represented as median and interquartile ranges. **(B)** Quantification of liver HIF1-a mRNA expression using qPCR of ad libitum fed and CR animals housed at different T_a_. Data are represented as median and interquartile ranges.**(C)** Quantification of liver mnSOD mRNA expression using qPCR of ad libitum fed and CR animals housed at different T_a_. Data are represented as median and interquartile ranges. *P<0.05, **P≤0.005, ****P≤0.0001.

**Figure 5.**
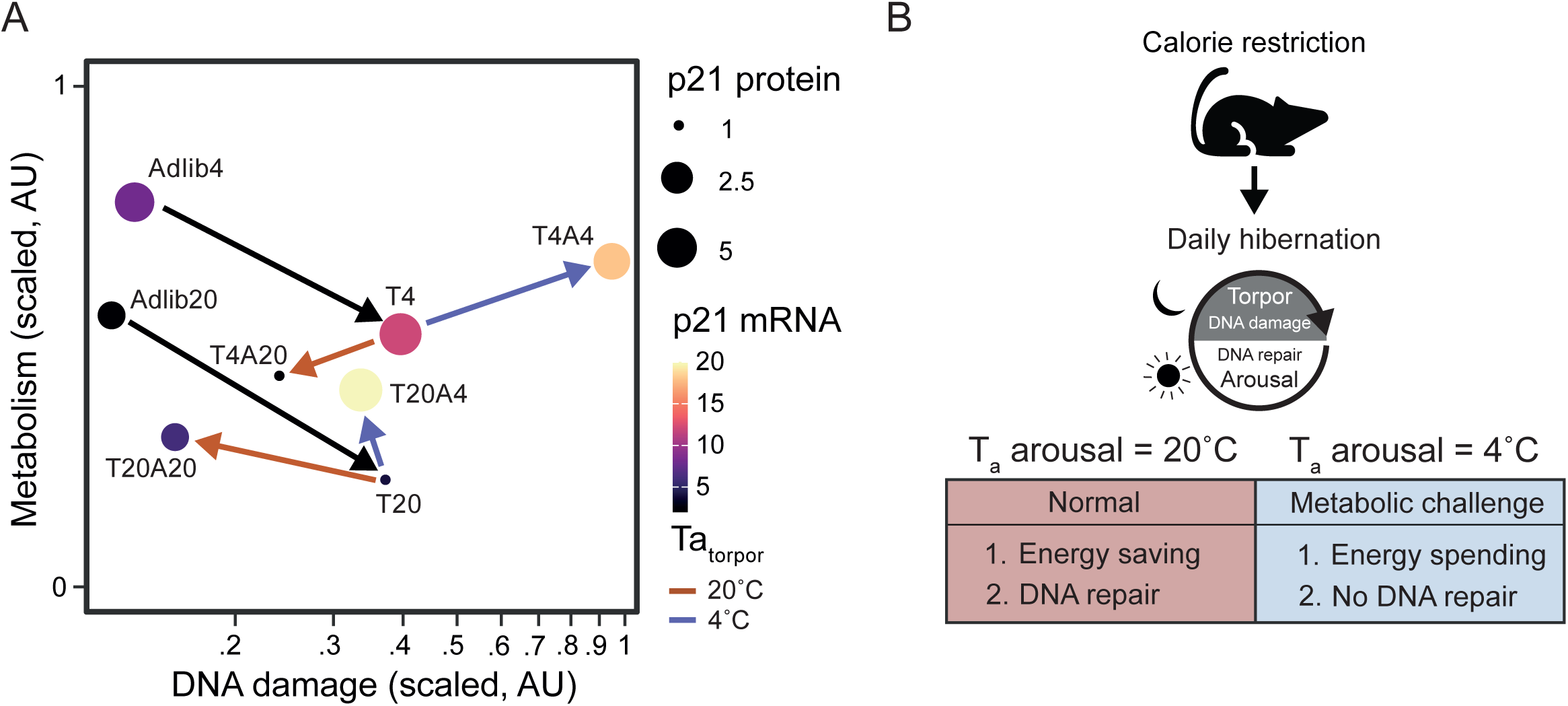
Metabolic Challenge Inhibits Repair of Torpor-induced DNA damage in Calorie Restricted Mouse. **(A)** Graphical representation of DNA damage, metabolic rate and p21 responses in ad libitum fed animals and CR animals housed at different T_a_. Entry into torpor (black arrows) yields similar metabolic rate reduction and DNA damage induction. Arousal at T_a_ of 20C (orange arrows) converge towards an optimum, with low metabolic rate compared to ad libitum fed animals, and DNA damage repair. Arousal at T_a_ of 4 degrees (blue arrows) increases metabolic rate further, and DNA damage is not repaired and even increased if preceded by a torpor at T_a_ of 4C. **(B)** Model of calorie restriction-induced daily torpor and arousal cycles, characterized by saving energy and repair of torpor-induced DNA damage at T_a_ of 20C, and characterized by spending energy and no repair or even induction of DNA damage at T_a_ of 4C.

## Discussion

The 30% calorie restriction regimen in mice housed at room temperature used in this study is a widely used experimental paradigm for inducing longevity. Yet, we here demonstrate this regimen to induce serial daily torpor during which DNA strand breaks accumulate in multiple tissues, as evidenced by comet assays of splenocytes, ɣH2AX foci in liver and kidney and accumulation of the downstream cell cycle inhibitor p21. Notably, DNA damage is fully resolved during the next euthermic period (‘arousal’) and repair is completed before the onset of the next torpor bout. Furthermore, an acute metabolic challenge induced by lowering T_a_ to 4°C during torpor does not further increase DNA damage, but markedly impairs DNA repair during arousal, suggesting that the DNA damage observed in CR originates from absence of repair of stochastic strand bread, rather than from induction of damage through oxidative stress. Collectively, these data demonstrate that CR mice experience repetitive daily cycles of DNA damage accumulation and repair, a phenomenon that may contribute to the lifespan-extending effects of calorie restriction.

Tissue analysis by comet assay and ɣH2AX staining clearly demonstrate DNA damage in multiple tissues of CR mice. To investigate the nature of DNA damage, we employed an acute metabolic challenge during specific phases of daily hibernation. Particularly, the metabolic challenge throughout torpor and arousal (T4A4) exacerbates excess DNA damage by inhibiting DNA repair, and increases p21 transcription and protein levels. Such incomplete DNA repair was also observed in mice exposed to cold solely during arousal (T20A4). In contrast, mice exposed to cold during torpor and subsequently returned to T_a_ of 20°C display the expected DNA repair during their arousal. Thus, metabolic challenge does not affect the induction of DNA damage during torpor but rather inhibits DNA repair during arousal. These data support a model in which inhibition of repair drives accumulation of DNA strand breaks in CR mice; in torpor repair is inhibited grace to the suppressed metabolism, whereas in arousal repair is suppressed when excess energy is needed to maintain body temperature. Given the upregulation of p21 in animals at low T_a_ during torpor and arousal (T4A4), our data also implies activation of DDR signaling during arousal in CR mice undergoing daily hibernation. Further, our data clearly demonstrates that in animals housed at 20°C, DNA repair is complete by the end of arousal, i.e. just prior to the next torpor bout, as evidenced by normalization of tail DNA, ɣH2AX foci and p21 expression. DNA damage and repair in CR mice are similar to those observed in seasonal hibernators, which recruit components of the DNA repair pathways during the hibernation season (17) and show an accumulation of DNA strand breaks during torpor bouts (10). Collectively, our data demonstrates that calorie restriction in mice induces a repetitive cycle of DNA damage and repair which is breached by a metabolic challenge.

Further, in T4A20, we observed that the torpor at 4°C precluded the subsequent arousal at 20°C to fully repair DNA damage. We speculate that the nature of the DNA damage animals suffer during a torpor bout at 4°C may be different from that at 20°C. We speculate torpor at 20°C may generate more simple lesions such as base lesions and SSBs, which are efficiently repaired using base excision repair during arousal. Conversely, 4°C torpor DNA damage may generate complex lesions, which require HR-based repair that is slower and potentially exceeds repair capacity of a single arousal bout. This remains to be experimentally validated in future experiments.

It is likely that mice entering an arousal at T_a_ of 4°C spend excess energy to rewarm T_b_ from ∼23°C to 36°C, which likely limits energy available for DNA repair. DNA repair is an energy-intensive process that requires ATP and deoxy nucleoside triphosphates (dNTPs) as building blocks to synthesize new DNA (18). During an arousal at 4°C, metabolism is likely redirected towards heat production, leading to a shortage of ATP, hampering the adequate energy supply and dNTP synthesis needed for effective DNA repair. However, this proposition requires that the organs in which DNA damage was assessed (liver, kidney and spleen) actively participate in heat production in animals housed in cold conditions. Although heat production during cold exposure is primarily attributed to brown adipose tissue (19), the liver, as a primary metabolic organ, also plays a crucial role in thermogenesis (20–22). Moreover, calorie restricted mice - be it housed at 20°C or 4°C - rely on thermogenesis in liver and other organs even more than *ad libitum* fed mice, as they lack significant fat reserves (23, 24). However, we have not directly assessed ATP, dNTP or NAD+ levels, which is a limitation.

As an alternative, DNA damage during daily hibernation may be caused by excess production of oxygen radicals. It has been proposed that increased or altered mitochondrial respiration under cold conditions (25) may elevate reactive oxygen species (ROS) production (12), which can lead to single-strand and double-strand DNA breaks (18). However, DNA damage has been reported to depend on increased nuclear ROS rather than mitochondrial ROS (26), which is in accordance with our results showing that increased metabolism during torpor does not increase the number of DNA strand breaks. Moreover, we observe that Nrf2 is decreased during torpor at both T_a_ of 20°C and 4°C without increasing DNA damage. Therefore, we postulate that the DNA damage observed during torpor is the result of stochastic DNA damage accumulation as a consequence of halted DNA repair due to the lowering of metabolism and/or T_b_. On the other hand, antioxidant gene expression is reduced during arousal even at T_a_ of 4°C, possibly as part of the transcriptional program associated with torpor and arousal. This would put arousing animals at risk of ROS during a metabolically active rewarming, increasing DNA damage as we observed during an arousal at T_a_ of 4°C.

A large body of literature documents the beneficial effects of CR in mice, including enhanced DNA repair capacity (27), longevity (28, 29), reduced senescence and inflammation (30) and a lowered incidence of degenerative diseases (31). Remarkably, CR extends life span twofold in an ERCC1^-/-^model of DNA repair deficiency (32) and to date, no single intervention that aims to mimic individual CR features has shown similar lifespan extension (33). Therefore, it is possible that the repetitive torpor and arousal cycles of the CR induced natural daily hibernation program in mice is responsible for the beneficial effects of CR. Part of this may hinge on a superior tissue repair in daily hibernation, similar to the effect daily hibernation in in the APP/PS1 Alzheimer’s mouse model, in which a single torpor and arousal sequence restored memory and hippocampal synaptic transmission (6). The increased longevity in CR mice might therefore result from daily hibernation, driving daily surveillance and repair of DNA damage promoting genomic integrity and thereby cellular and tissue health. Whether torpor-associated DNA repair cycles causally contribute to CR-mediated lifespan extension remains to be established experimentally.

## Materials and Methods

### Animals

Adult male and female C57BL/6J mice were bred in the institutional facility for animal studies at the University Medical Center Groningen (CDP-UMCG). Ethical clearance was obtained from the Central Committee on Animal Experiments of The Netherlands (CCD license number AVD10500202010706) and animal welfare body UMCG (protocol 2020706-01-002). Mice were housed individually under a 12h light-dark (L:D) cycle and fed with standard chow (Ssniff V1554-703). After one week of acclimatization, a temperature logger (G2 E-Mitter transponders, Starr Life Sciences, Oakmont, PA) was implanted intra-abdominally under isoflurane anesthesia. CR mice were subjected to 30% calorie restriction, which was achieved by gradual reduction from 0 to 30% restriction within 2 weeks with free access to water. Control animals were provided food ad libitum. The mice were divided into eight experimental groups: Adlib20, Adlib4, T20, T4, T20A20, T20A4, T4A20, and T4A4 (Fig. 1A). Mice were euthanized by exsanguination under isoflurane anesthesia. The spleen was immediately harvested, and splenocytes were obtained using a 10 µm mesh in ice-cold phosphate-buffered saline (PBS). Other tissues were harvested after the mice were flushed with saline 0.9% for 15 min by snap freezing in liquid nitrogen and stored at -80°C.

### Calorimetric Cages

After at least four weeks of calorie restriction and stable body weight, mice were placed into calorimetric cages in climate-controlled cabinets (indirect calorimetry PhenoMaster, TSE systems, Germany), to measure body temperature, oxygen consumption, carbon dioxide production and activity during the final 2 days of the experiment, during which mice were exposed to 20°C and/or 4°C (Fig. 1A). Mice exposed to 4°C during arousal were given additional food to compensate for the increased energy expenditure at low T_a_.

### Comet assay

A comet assay was performed on freshly isolated splenocytes, according to the manufacturer’s instructions (Trevigen, USA). Alkaline comets were used to assess DNA for both single- and double-strand breaks, while neutral comets assessed only double-strand breaks. Cell suspension containing 5-10 x 10^6 cells/mL was mixed at a 1:10 ratio with low melting agarose, then pipetted onto comet slides. Electrophoresis was run at 21V for 30 minutes. After washing, the slides were dried at 37°C for 30 min and stained with SYBR GOLD. Images were captured using a Leica DM 2000 LED epifluorescence microscope at 10x magnification. Quantification was performed using ImageJ, using the OpenComet plugin combined with manual quantification using the Comet Assay plugin. DNA damage was evaluated based on the percentage of DNA intensity in the comet tail (%Tail DNA). A minimum of 100 cells per animal was quantified.

### Immunofluorescence staining

Cryosections of liver and kidney (8 µm) were fixed with 3% paraformaldehyde and washed with PBS. Permeabilization was performed using 0.1% Triton X-100 for 15 min, followed by washing with PBS. Slides were incubated overnight at 4°C with anti-histone H2AX (p-Ser139; Novus NB100-384) in 1% bovine serum albumin (BSA; Capricorn Scientific, Germany). After washing, the slides were incubated with a secondary antibody (Alexa Fluor 594, Thermo Fisher) for 1h at RT. DAPI was used to visualize nuclei, and the slides were mounted with Vectashield (Vector Laboratories, USA). Images were captured using a Leica SP8 confocal microscope. Quantification was performed by counting foci per nuclei of at least 50 cells per animal, using Imaris software 9.7.2 (Oxford Instruments).

### P21 RNAscope

Frozen liver (4 µm) and kidney (8 µm) sections were mounted on poly L-lysine-coated glass slides (Epredia SuperfrostTM Plus; USA). The tissues were fixed with 4% paraformaldehyde at 4°C for 15 min. Multiplex in-situ hybridization was performed using the RNAscope Multiplex Fluorescent Reagent Kit v2 (ACD, Biotechne, USA) according to the manufacturer’s instructions. After counterstaining with RNAscope DAPI for 1 min at RT, the slides were mounted with VECTASHIELD (Vector Laboratories, USA) and dried for 30 min in the dark. Images were captured using a Leica DM 2000 LED epifluorescence microscope or Leica Thunder at 20x magnification. Quantification was performed using ImageJ software.

### Quantitative PCR

RNA was isolated from liver tissue using a commercial kit (Macherey-Nagel), following the manufacturer’s instructions. RNA purity and concentration were determined using a UV spectrophotometer (NanoDrop ND 1000; Thermo Fisher Scientific, USA). cDNA synthesis was performed using a reverse transcriptase mix according to the manufacturer’s instructions (Promega). Twelve random samples were pooled to generate a standard curve. For transcript analysis, cDNA was amplified on Biorad CFX 384 using Promega GoTAQ master mix (Promega, Madison, USA) using the desired primer. Quantification was performed by calculating the delta Ct for a housekeeping gene (GAPDH or β-actin). Primer sequences of genes targets can be found in SI appendix table 2.

**Table 2.**
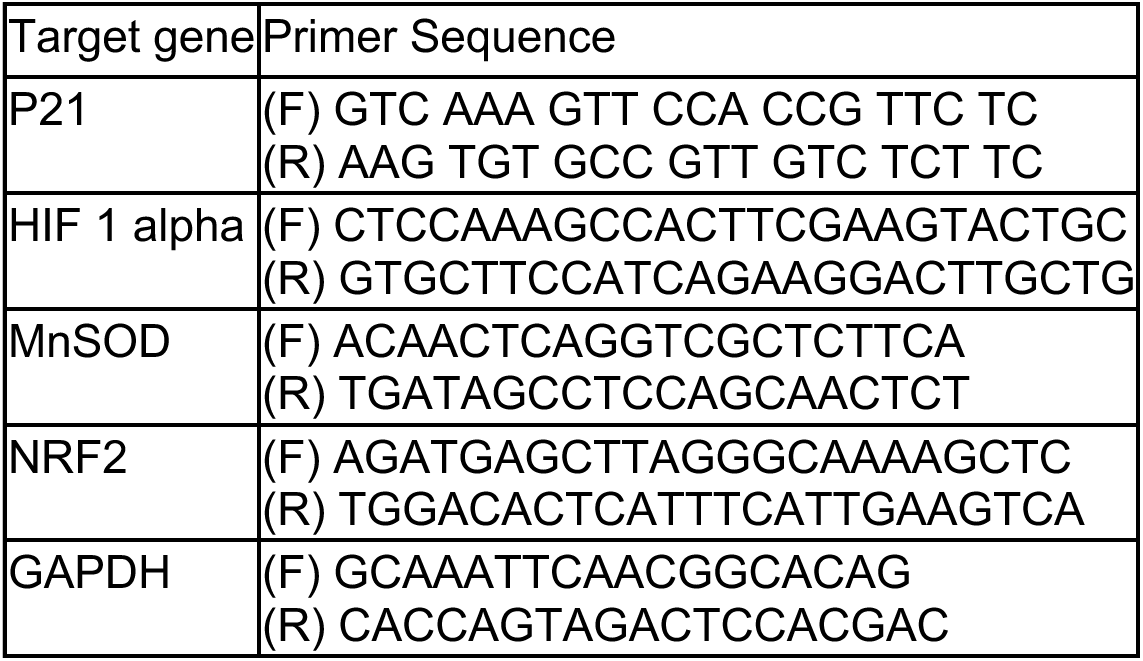
Primer sequence of target genes.

### Western Blot

Frozen liver tissue (50 mg) was homogenized using a minithorax in 500uL RIPA T_b_S (50mM Tris-HCl, 150mM NaCl, 1% Igepal Ca-630, 1%, 0.5% sodium deoxycholate, 1% SDS) supplemented with inhibitors (40uL/mL protese inhibitor cocktail, 10uL/mL sodium orthovanadate, 10uL/mL sodium fluoride and 0.71uL/mL beta mercaptoethanol). Lysates were incubated on ice for 30 min and centrifuged at 14,000 × g for 20 min at 4°C. The supernatant was collected and the protein concentration was measured using a Bio-Rad protein assay kit. A total of 20 µg was loaded into a TGX stain-free gel (Bio-Rad, USA) the electrophoresis run using a Mini Protean Tetracell system (Bio-Rad) at 200V for 30 min. Following electrophoresis, proteins were transferred to a nitrocellulose membrane (Bio-Rad) at 2.5A, 25V for 7 min using a Trans-blot Turbo transfer system (Bio-Rad) in Turbo blot buffer. Membranes were blocked with 5% skim milk (Sigma Aldrich, USA) in

Tris-buffered saline containing 0.1% Tween-20 (T_b_S-T) for 20 min at RT. After blocking, the membranes were incubated overnight at 4°C with primary antibodies anti-p21 (1:1000; Abcam) diluted in 3% BSA in T_b_S-T. After washing, the membranes were incubated with goat anti-rabbit IgG/HRP (DAKO Agilent, Santa Clara, USA) for 1h at RT. Protein bands were detected using Western Lighting Ultra (Revvity, Massachusetts, USA) and visualized with the ChemiDoc MP Imaging System (Bio-Rad). Band intensities were quantified using the Image Lab software (Bio-Rad) with total protein correction.

### Statistical analysis

Statistical analysis were performed using GraphPad Prism (GraphPad Prism software 9.1.0 GraphPad Windows, San Diego, California, USA) and R version 4.5.1 (2025-06-13). For statistical analysis, one-way analysis of variance (ANOVA) with Tukey’s post-hoc analysis was used for normally distributed data. Kruskal-Wallis test was used for skewed data, with Dunn’s multiple comparison test as a post-hoc test. Statistical significance was set at P < 0.05. A cyclic generalized additive model was fit to 3 days of measurements of body temperature and metabolism using the gam function from package mgcv for ad libitum and CR fed animals.

## Acknowledgements

FFL was supported by LPDP scholarship (The Indonesian Endowment Fund for Education, Ministry of Finance of the Republic of Indonesia) for her PhD trajectory at University of Groningen [contract number 2020032230760]. The funder had no role in study design, data collection and analysis, or writing of the manuscript.

---

**Supplementary Figure 1.**
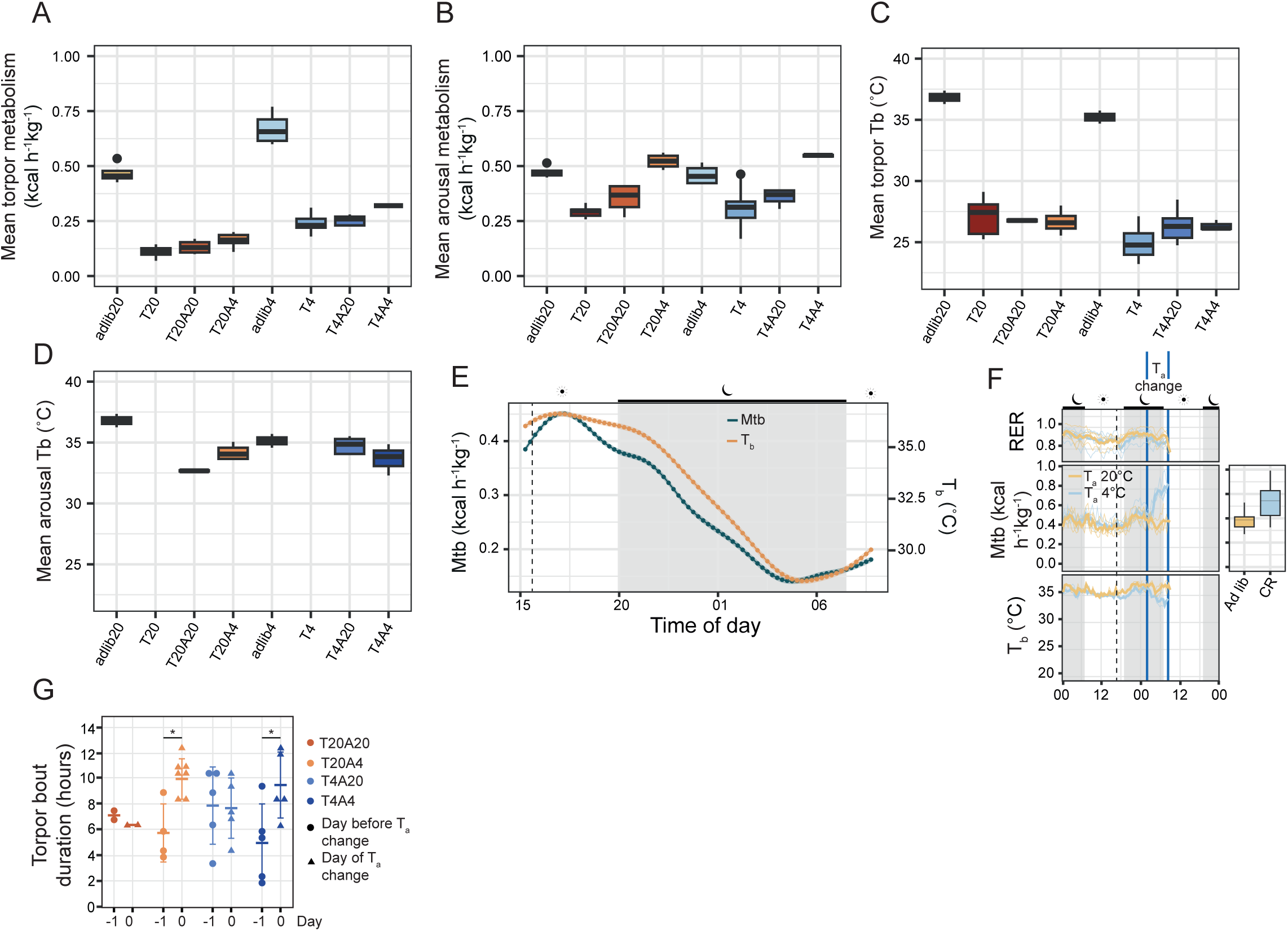
CR induces changes in metabolic rate, body temperature, RER, which are influenced by ambient temperature. **(A)** Mean metabolic rate per animal measured during torpor for CR animals, and during the same time frame for ad libitum fed animals. Data are represented as median and interquartile ranges. **(B)** Mean metabolic rate per animal measured during arousal and during the same time frame for ad libitum fed animals. Data are represented as median and interquartile ranges. **(C)** Mean T_b_ per animal measured during torpor and during the same time frame for ad libitum fed animals. Data are represented as median and interquartile ranges. **(D)** Mean T_b_ per animal measured during arousal and during the same time frame for ad libitum fed animals. Data are represented as median and interquartile ranges. **(E)** Cyclic fits from multiple days of metabolic rate and T_b_ for CR animals entering torpor and initiating arousal. Data are represented as mean ± 95% CI **(F)** RER, metabolic rate and T_b_ of ad libitum fed animals exposed to T_a_ of 20°C and 4°C between 02:00 and 08:00. Data are represented as median and interquartile ranges. **(G)** Torpor bout duration measured on the day before changing T_a_, and on the day T_a_ was changed for CR animals. Data are represented as median ± SD. *P<0.05, **P≤0.005, ****P≤0.0001.

**Supplementary Figure 2.**
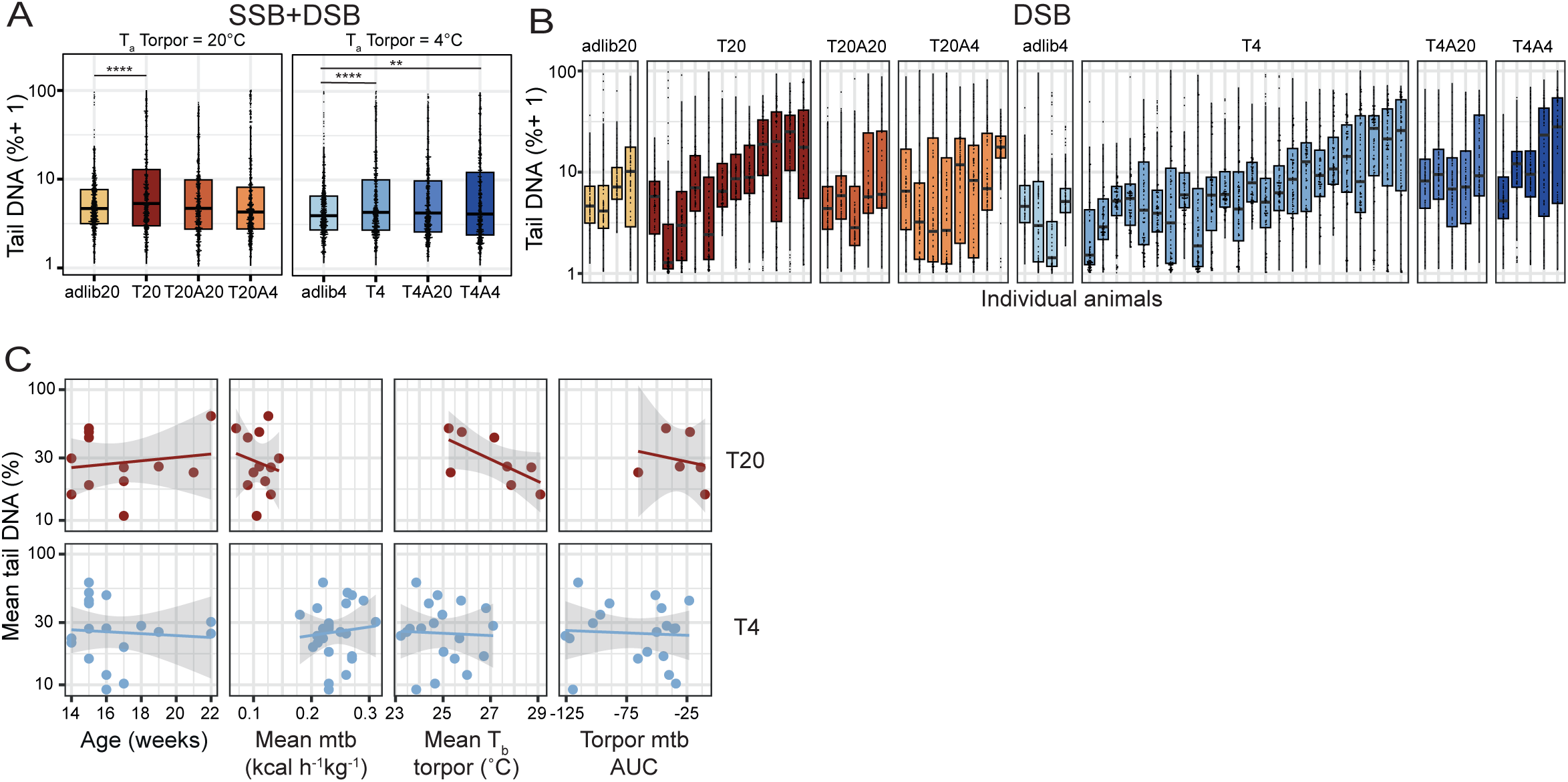
CR induces daily cycles of DNA damage and repair, with considerable variance between animals. **(A)** Quantification of alkaline comets (SSB+DSB) of ad libitum fed and CR animals housed at different T_a_. Data are represented as median and interquartile ranges. **(B)** Per-animal quantification of neutral comets of ad libitum fed and CR animals housed at different T_a_. Data are represented as median and interquartile ranges. **(C)** Relationships between mean tail % DNA, representing DNA damage, and age of the animals, mean metabolic rate during torpor, mean T_b_ during torpor and torpor metabolism AUC during torpor at T_a_ of 20C and 4C. *P<0.05, **P≤0.005, ****P≤0.0001.

